# The Alzheimer’s disease-associated protective *Plcγ2*-P522R variant promotes beneficial microglial functions

**DOI:** 10.1101/2020.04.08.031567

**Authors:** Mari Takalo, Rebekka Wittrahm, Benedikt Wefers, Samira Parhizkar, Kimmo Jokivarsi, Teemu Kuulasmaa, Petra Mäkinen, Henna Martiskainen, Wolfgang Wurst, Xianyuan Xiang, Mikael Marttinen, Pekka Poutiainen, Annakaisa Haapasalo, Mikko Hiltunen, Christian Haass

**Author notes:** These authors contributed equally (Mikko Hiltunen, Christian Haass).

## Abstract

**Background:** Microglia-specific genetic variants are enriched in several neurodegenerative diseases, including Alzheimer’s disease (AD), implicating a central role for alterations of the innate immune system in the disease etiology. A rare coding variant in the *PLCG2* gene (rs72824905, p.P522R) selectively expressed in microglia and macrophages was recently identified and shown to reduce the risk for AD.

**Methods:** To assess the role of this variant in the context of immune cell functions, we generated a Plcγ2-P522R knock-in (KI) mouse model using CRISPR/Cas9 gene editing.

**Results:** Functional analyses of macrophages derived from homozygous KI mice and wild type (WT) littermates revealed that the P522R variant potentiates the primary function of Plcγ2 as a Pip2-metabolizing enzyme. This was associated with improved survival, enhanced phagocytic activity, and increased acute inflammatory response of the KI cells. Enhanced phagocytosis was also observed in mouse BV2 microglia-like cells overexpressing human PLCγ2-P522R, but not in PLCγ2-WT expressing cells. Furthermore, the brain mRNA signature together with microglia-specific PET imaging indicated microglia activation in Plcγ2-P522R KI mice.

**Conclusion:** Thus, we have delineated cellular mechanisms of the protective Plcγ2-P522R variant, which provide further support for the emerging idea that activated microglia exert protective functions in AD.

## Findings

Recent genome-wide association studies have identified several Alzheimer’s disease (AD)-associated risk loci in genes selectively or preferentially expressed in microglia (e.g. *TREM2, ABI3, PLCG2*) [1]. A rare coding variant in microglia-/macrophage-specific *PLCG2* gene (rs72824905, p.P522R, OR<0.6) encoding phospholipase C gamma 2 (Plcγ2) enzyme was recently identified, showing a reduced risk of AD [1]. Interestingly, the same *PLCG2* variant associated with a lower risk of other neurodegenerative diseases and increased the likelihood for longevity [2]. Plcγ2 catalyzes the conversion of phosphatidylinositol 4,5-bisphosphate (Pip2) to inositol 1,4,5-trisphosphate (Ip3) and diacylglycerol (Dag) upon activation of various transmembrane immune receptors, including Triggering receptor expressed on myeloid cells 2 (Trem2) (**Fig 1A**) [3, 4]. Ip3 and Dag regulate pathways related to e.g. survival, phagocytosis, and cytokine production via controlling intracellular calcium mobilization as well as protein kinase C, nuclear factor kappa-light-chain-enhancer of activated B cells (Nfκb), mitogen-activated protein kinase (Mapk/Erk), and protein kinase B (Akt) signaling [5-7]. The P522R variation locates in the regulatory domain of Plcγ2, but whether this variant affects the above-mentioned functions of microglia and other myeloid lineage cells, is not known.

**Figure 1.**
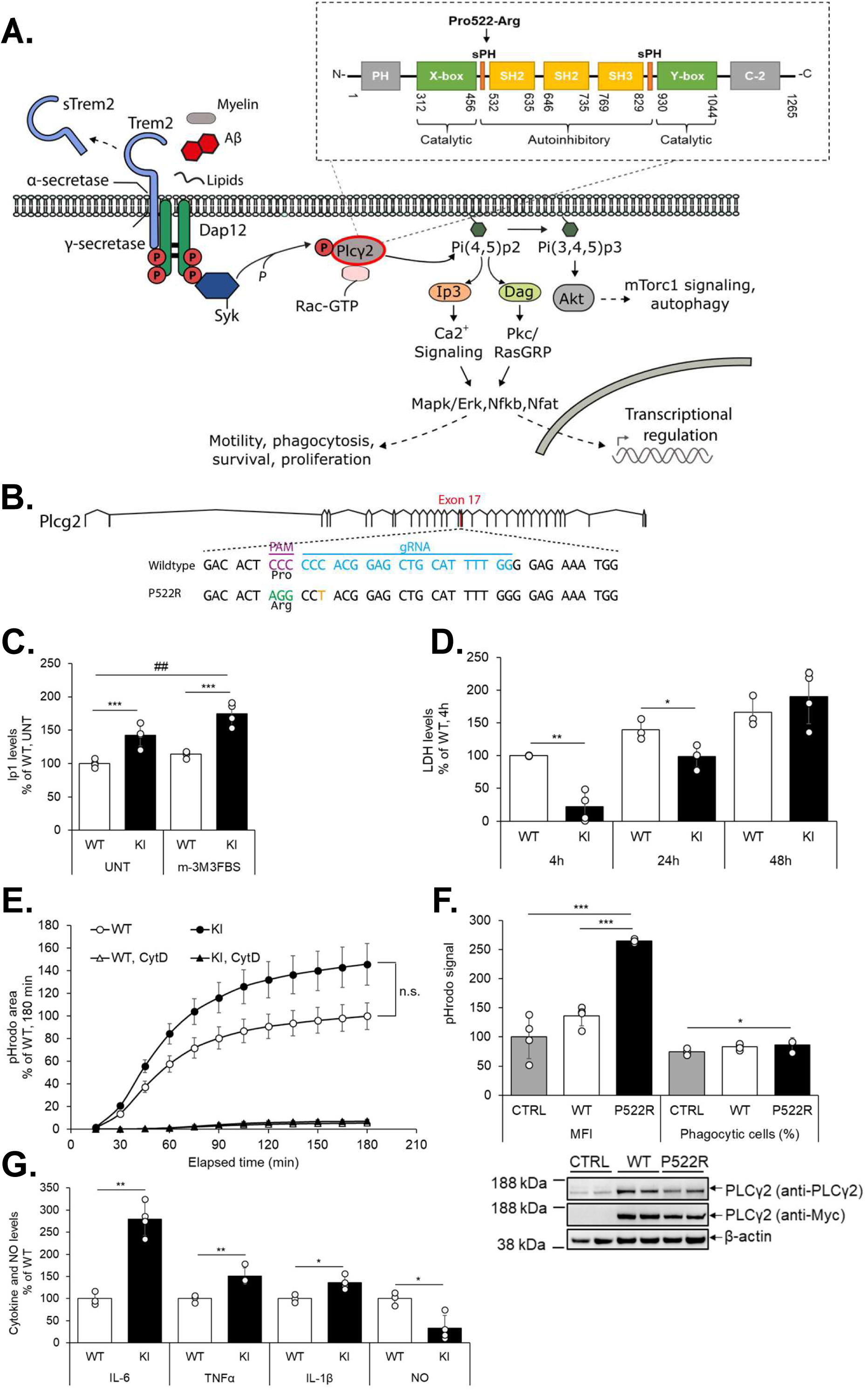
The Plcγ2*-*P522R variant increases Plcγ2 enzymatic activity, survival, phagocytic activity, and inflammatory response in macrophages. A) A schematic view of Plcγ2 signaling. B) Strategy to target murine *Plc*γ*2* locus, indicating protospacer region (blue), protospacer adjacent region (PAM, purple), and introduced nucleotide changes (missense: green; silent: orange). C) Inositol monophosphate (Ip1) formation in Plcγ2-P522R knock-in (KI) and wild type (WT) bone marrow-derived macrophages (BMDMs) in basal state and after 1h m-3M3FBS (phospholipase C agonist, 25µM) stimulation. Values are normalized to the total protein concentration. Mean ± SD, % of untreated (UNT) WT, n=4 per genotype, 2 technical replicates. Two-way ANOVA, LSD post hoc test, ***p<0.001 (genotype effect), ##p<0.001 (treatment effect). D) Lactate dehydrogenase (LDH) levels in KI and WT BMDMs after 4h, 24h and 48h macrophage colony stimulation factor 1 (mCSF) withdrawal. Values are normalized to the maximum LDH release within each well. Mean ± SD, % of WT (4h), n=3-4 per genotype, 2 technical replicates. Two-way ANOVA, LSD post hoc test, *p<0.05, **p<0.01. E) Phagocytic activity in KI and WT BMDMs. Cytochalasin D (CytD, 5µM) was used as a control. pHrodo signal is normalized to the cell count in each well. Mean ± SEM, n=6-7 per genotype, 3 technical replicates. Independent samples *t*-test, n.s. non-significant. F) Phagocytic activity in BV2 microglial cells overexpressing Myc-tagged human PLCγ2-P522R, PLCγ2-WT, or control (CTRL) vector shown as median fluorescence intensity (MFI, left panel) and percentage of phagocytic cells (right panel). Western blot (lower panel) confirming overexpression of P522R and WT constructs. Mean ± SD, % of CTRL, n=4. Mann-Whitney U test, *p<0.05, ***p<0.001. G) Interleukin-6 (IL-6), tumor necrosis factor α (TNFα), IL-1β, and nitric oxide (NO) levels in conditioned media of KI and WT BMDMs upon 3h lipopolysaccharide (LPS) and interferon gamma (IFNγ)-treatment. Values are normalized to the total protein concentration in the corresponding wells. Mean ± SD, % of WT, n=3-4 per genotype. Independent samples *t*-test, *p<0.05, **p<0.01.

Here, we have generated a Plcγ2-P522R knock-in (KI) mouse model using the CRISPR/Cas9 gene editing technology (**Fig 1B**) [8, 9] to investigate the effects of this protective variant in isolated macrophages and microglia and *in vivo* within the brain. Founder mice with successful incorporation of the variant and confirmed to be negative for off-target effects were used for generation of a viable and fertile homozygous Plcγ2-P522R line (**Fig. S1**).

### The P522R variant increases Plcγ2 enzyme activity and enhances survival, phagocytosis, and inflammatory response

Bone marrow-derived macrophage (BMDM) cultures were established from *femur* and *tibia* bones of six-month-old Plcγ2-P522R KI mice and their wild type (WT) littermates [10-12]. The P522R variant has previously been shown to increase the production of inositol phosphate (Ip1, a surrogate of Ip3) and intracellular calcium release in transiently transfected HEK293 and COS cells [13]. To assess Plcγ2 enzyme activity in KI myeloid cells, Ip1 formation was measured from BMDM lysates as previously described [6, 13]. A significantly higher basal Ip1 levels was detected in cultured KI BMDMs as compared to WT cells, and treatment with the phospholipase C agonist m-3M3FBS [7] further increased the levels of Ip1 in both, KI and WT cells (**Fig. 1C**). Thus, these findings indicate that the P522R variant exerts a hypermorphic effect on the basal enzyme activity of Plcγ2 in cultured mouse macrophages.

Next, we examined the functional consequences of increased Plcγ2 activity in cultured BMDMs. A cytotoxicity assay revealed significantly lower levels of lactate dehydrogenase (LDH) in the conditioned medium of KI BMDMs as compared to WT cells after withdrawal of macrophage colony stimulation factor (m-CSF) (**Fig 1D**), suggesting that the protective variant mitigates entering of BMDMs to an apoptotic state. Importantly, Trem2 deficiency has been shown to compromise the survival of microglia and peripheral macrophages [12-15], whereas antibody-mediated stabilization of mature Trem2 strongly enhances the survival and proliferation of BMDMs (Schlepckow et al., EMBO Mol Med, in press) [16]. This supports the idea that Trem2 and Plcγ2 act on the same pathway (see **Fig. 1A**), and that the Trem2 risk and Plcγ2 protective variants have opposite effects on cell fate.

Activation of the Trem2 pathway is also known to enhance phagocytosis, a beneficial cellular process removing β-amyloid, apoptotic neurons, and other pathological substances in the aging or diseased brain [11, 12, 17, 18]. Here, imaging of pHrodo-labeled bioparticle uptake by BMDMs showed a trend towards higher phagocytic activity in KI as compared to WT cells (**Fig 1E**). To confirm that the protective variant enhances phagocytosis, cDNAs encoding human PLCγ2-WT and PLCγ2-P522R were expressed in microglia-like BV2-cells. Significantly higher fluorescence intensity was detected in BV2-cells overexpressing the P522R variant as compared to those overexpressing PLCγ2-WT or a control plasmid (**Fig 1F**). Simultaneously, the overall percentage of phagocytic cells was only marginally higher in BV2-cells overexpressing the PLCγ2-P522R variant, suggesting that the phagocytic capacity per cell was increased.

Plcγ2 has been suggested to promote Nfκb-mediated innate immune response and cytokine production in peripheral immune cells [19]. To assess an acute response of the BMDMs to an inflammatory stimulus, the cells were treated with lipopolysaccharide (LPS) and interferon-gamma (IFNγ) for three hours. BMDMs from KI mice released significantly higher levels of tumor necrosis factor-α, (TNFα), interleukin-6 (IL-6), and −1β (IL-1β) into the culture medium (**Fig. 1G**), suggesting that macrophages expressing the P522R variant display stronger and/or faster response to the stimulus. Despite higher cytokine secretion, significantly lower levels of nitric oxide (NO) were detected in the culture medium of KI BMDMs. Importantly, the IFNγ-induced production of NO exacerbates apoptosis and compromises cell viability in a variety of cell types. Furthermore, the neurotoxic effect of activated microglia is largely mediated by NO [20-22]. Thus, our observation supports the idea that the P522R variant protects cells upon different stress conditions.

### The mRNA profile associated with Plcγ2-P522R suggests activation of microglia and modulation of Plcγ2 signaling

Next, we searched for potential molecular changes in the brain of six-month old KI and WT male mice. Changes in total brain mRNA expression were determined utilizing a Nanostring neuropathology gene expression panel. Out of 770 analyzed genes, 57 (7%) were significantly up- and 32 (4%) significantly downregulated in the brain of KI as compared to WT mice (**Fig. 2A**). Notably, several genes directly linked to Plcγ2 signaling (e.g. *Itpr1, Camk2d, Mapk3/Erk1, Rac1*, and *Rhoa*) showed a significantly elevated expression in the KI mouse brain (**Fig 2B**). *Itpr1* encodes an Ip3 receptor and acts directly downstream of Plcγ2. Similarly, Mapk/Erk-signaling is induced by Plcγ2 activation [6] and regulates pathways related to survival, proliferation, differentiation, and inflammatory responses in brain immune cells and other cell types [23]. Rac1 and RhoA can be activated in a Plcγ2 and/or Mapk/Erk-dependent manner [24-25] and play pivotal roles in microglia activation, migration, phagocytosis, and neuronal loss [27-29]. Thus, the higher expression of such targets could be directly associated with Plcγ2-dependent Ip3 signaling and is in line with improved survival, inflammatory response, and phagocytosis observed in KI macrophages.

**Figure 2.**
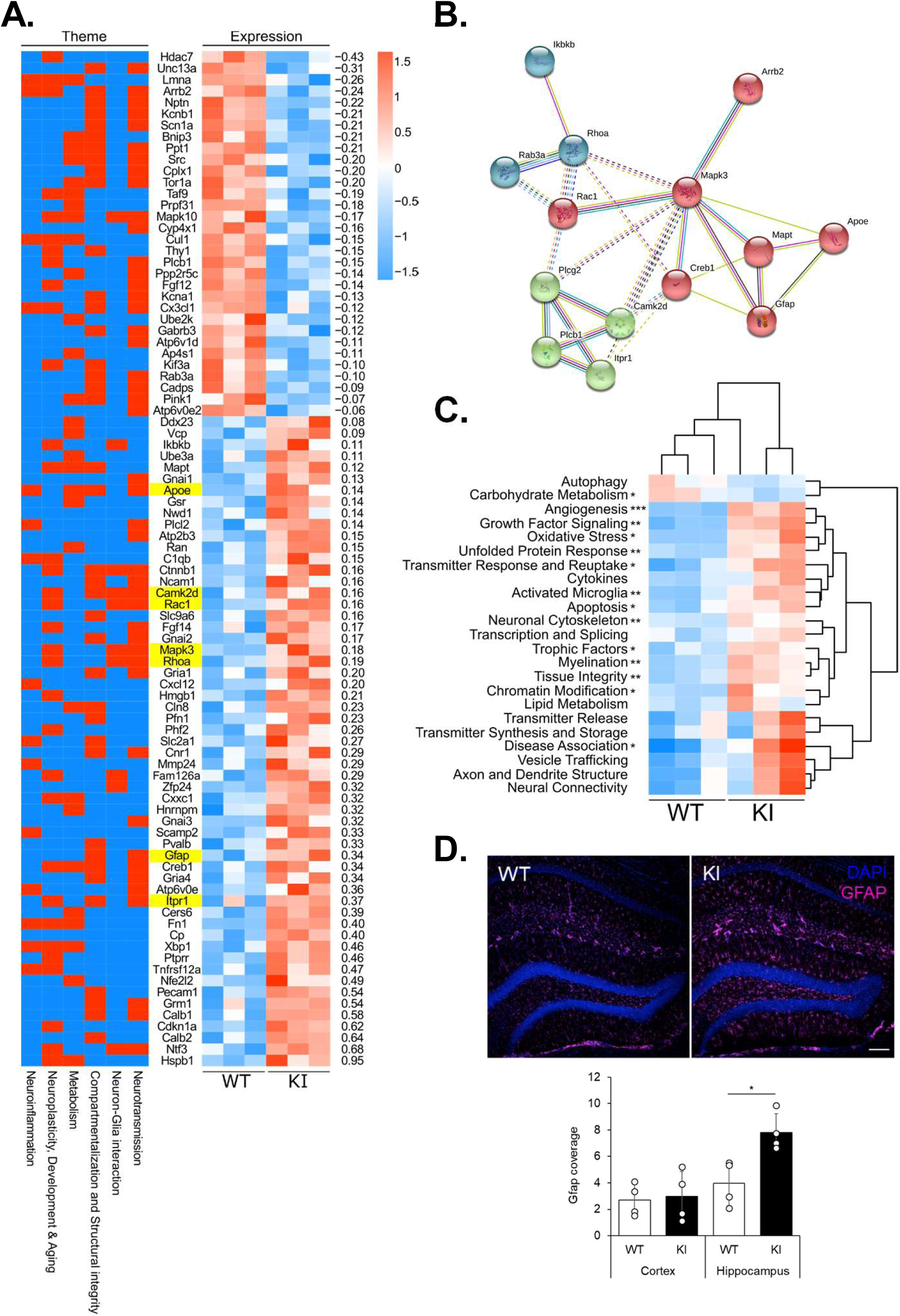
Activated microglia and astroglia in brains of mice expressing the *Plcγ2-P522R* protective mutation. A) A heatmap showing significantly (p<0.05) changed gene expression and associated themes obtained from the neuropathology panel. Targets are arranged according to log2-transformed fold-changes in Plcγ2-P522R homozygote knock-in (KI) mice as compared to the wild type (WT) littermates, n=3 per genotype. B) A String-network graphic and kmeans clustering of significantly affected targets associated with *Plcg2* (https://string-db.org/cgi/network.pl?taskId=AF3kTOpi6ovG). C) A heatmap showing pathway scores for neuropathology panel-derived pathway annotations. Significantly changed pathways are indicated with asterisk, n=3 per genotype, Independent samples *t*-test, *p<0.05, **p<0.01, ***p<0.001. D) Representative images of Gfap-stained hippocampus from WT and KI mice (upper panel). Significantly higher levels of Gfap were detected in the hippocampus, but not in the cortex of KI mice (lower panel). Scale bars represent 10μm. Mean ± SD, n=4 per genotype, two-tailed unpaired *t*-test, *p<0.05.

Pathway analysis demonstrated a significant upregulation of microglia activation among several other biological pathways in the brain of KI as compared to the WT mice (**Fig 2C**). Recent transcriptomic studies in CNS immune cells have identified a unique subtype of disease-associated microglia (DAM), which is characterized by high expression of pro-survival, lipid metabolism, and phagocytosis-associated genes [30, 31]. Activation of DAM is driven in a Trem2-dependent manner and it is essential for impeding AD-related β-amyloid pathology [31]. To assess whether the protective variant shifts microglia towards the DAM phenotype, expression of DAM signature genes was examined. A significant upregulation of *Apoe*, which is one of the key upregulated DAM genes in microglia [30-32], was detected in the brain of KI mice (**Fig. 2A**), while other targets, such as *Trem2, Itgax*, and *Cd68*, remained unchanged. Additional qPCR-based analyses of microglia-specific DAM genes showed a trend towards increased expression of *Cst7, Tyorobp, Clec7a*, and *Ccl3* in the brain of KI mice (**Fig. S2**). Moreover, a significant upregulation of an astrocytic marker, *Gfap*, was found in the brain of KI mice (**Fig. 2A**). Immunohistochemical analysis revealed a significantly stronger staining of Gfap and a hypertrophic morphology of astrocytes in the hippocampus of KI as compared to WT mice (**Fig. 2D**). This is an interesting finding also in the context of Plcγ2-P522R-associated RNA profile as *Apoe* is primarily expressed in the astrocytes. Collectively, these results suggest a potentially beneficial crosstalk between activated microglia and reactive astrocytes in the hippocampus of Plcγ2-P522R KI mice.

### The Plcγ2-P522R protective variant increases microglia activation in the brain of KI mice

Finally, we elucidated the activation state of microglia in the Plcγ2-P522R KI mice *in vivo* by conducting PET imaging using the 18F-FEPPA radioligand specific for the translocator protein of 18 kDa [33, 34]. A significant increase in 18F-FEPPA signal was detected in KI compared to WT mice in all analyzed brain areas in one-year-old female mice (**Fig. 3A**). Interestingly, reduced microglia activation has been observed in mice carrying *Trem2* loss-of-function mutations [12, 35, 36], suggesting that the protective Plcγ2*-*P522R variant and the risk-increasing *Trem2* variants exert opposite effects on microglia function. Furthermore, impaired microglia activation has been shown to correlate with reduced cerebral glucose metabolism. However, microglial hyperactivation in *Grn* knockouts also leads to reduced cerebral glucose metabolism [12,36] demonstrating that the two extremes of microglial activation states are both deleterious. To prove, if the microglial activation triggered by the Plcγ2*-*P522R variant affects glucose metabolism, we measured cerebral metabolic rate of glucose using 18F-FDG-PET [33]. FDG-PET did not detect significant differences in any of the brain regions studied nor any correlation between 18F-FEPPA and 18F-FDG signals supporting the idea that the Plcγ2*-*P522R variant supports a protective microglial activation state.

**Figure 3.**
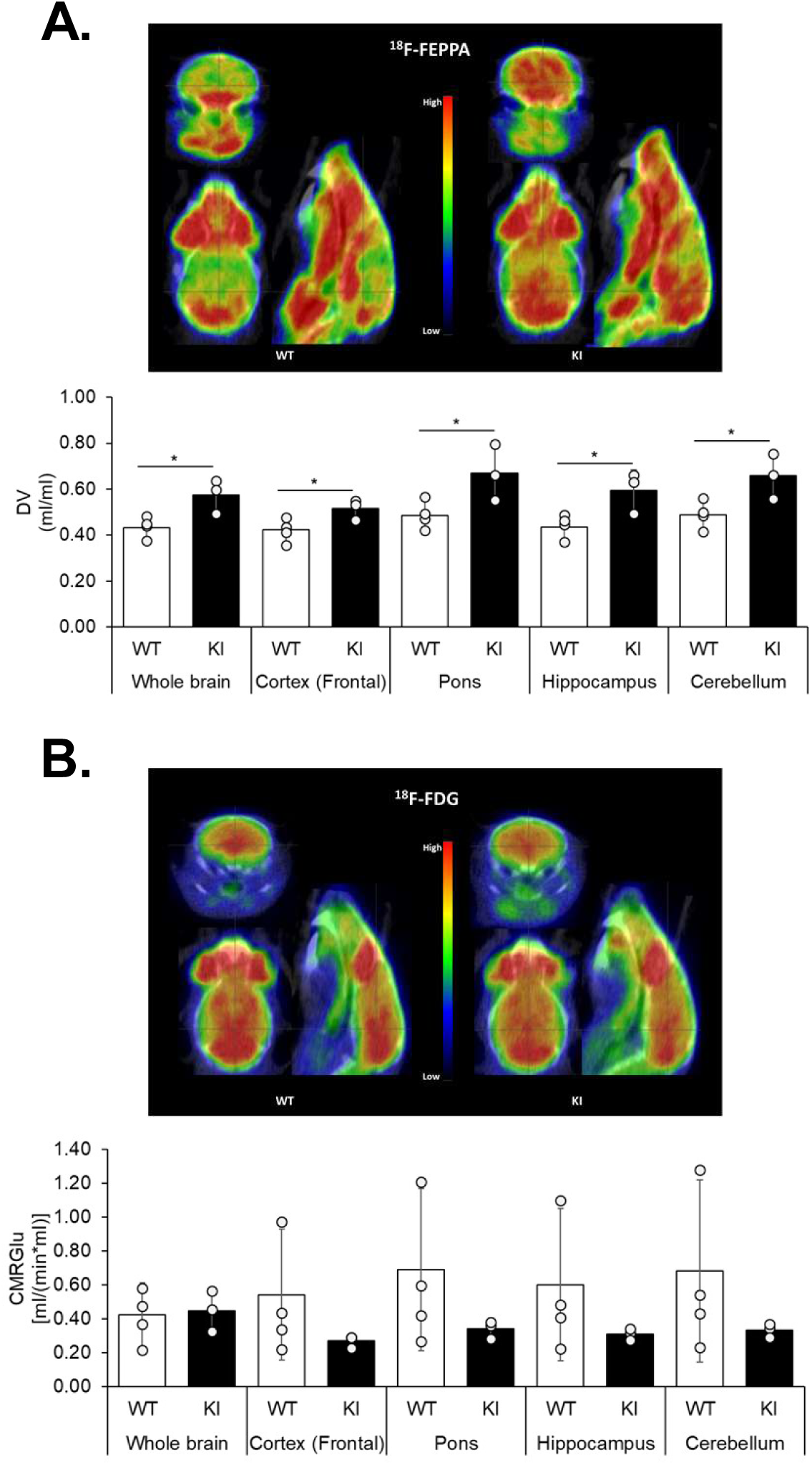
Plcγ2-P522R expression increases microglia activation in the brain of one-year-old mice. A) Representative PET-images of the uptake of 18F-FEPPA (time window 15-70min) (upper panel) in the brains of Plcγ2-P522R homozygous knock-in (KI) and wild type (WT) female mice at one year of age. Estimated total distribution volume (DV, ml/ml) in total brain, frontal cortex, pons, hippocampus, and cerebellum calculated with the Logan model using image derived input function (IDIF, lower panel), n=3-4 per genotype. Independent samples *t*-test, *p<0.05. B) Representative PET images of the uptake of 18F-FDG (time window 20-80min) in WT and KI mouse (upper panel). Cerebral metabolic rate of glucose [MRGlu, (ml/min*ml)] was estimated using the Patlak model (with lumped constant of 0.71, lower panel), n=3-4 per genotype. Independent samples *t*-test.

In conclusion, our results suggest that the protective Plcγ2-P522R variant potentiates the primary function of Plcγ2 and enhances important immune cell functions *in vitro* and microglia activation *in vivo*. Although further studies focusing on the microglia-specific effects of the Plcγ2-P522R variant upon AD-associated stress are required, the present findings may indicate protective pre-priming of microglia, which could be beneficial in the brain in neurodegenerative conditions.

## METHODS

### Animals

Plcγ2-P522R KI mice were generated by CRISPR/Cas9-assisted gene editing in mouse zygotes as described previously [8, 9, 12]. Briefly, pronuclear stage zygotes were obtained by mating C57BL/6J males with superovulated C57BL/6J females (Charles River). Embryos were then microinjected into the male pronucleus with an injection mix containing *Plc*γ*2*-specific CRISPR/Cas9 ribonucleoprotein (RNP) complexes. RNPs consisted of 50 ng/µl S.p. Cas9 HiFi protein (IDT), 0.6 µM crRNA (protospacer CCAAAATGCAGCTCCGTGGG; IDT), 0.6 µM tracrRNA (IDT), and 25 ng/µl mutagenic single-stranded oligodeoxynucleotide (ssODN) (5′ - CGCACTGGTCCTACTCTCCACCTTCTTGTGGAACCATTTCTCCCCAAAATGCAGCTCCGTAGGCCTAGTGTCCTGAGCCACAAGCATCCGAAAGGGCTTATTACAGCTCGCTCTGCCCTCT CCTGA□3′), comprising the P522R substitution and an additional silent mutation for genotyping purposes (IDT; see also Fig 1B). After microinjection, zygotes were cultured in KSOM medium until they were transferred into pseudopregnant CD-1 foster animals. To identify putative off-target sites of the *Plc*γ*2*-specific guideRNA, the online tool CRISPOR (http://crispor.tefor.net/) was used. For analysis, genomic DNA of WT and heterozygous Plcγ2-P522R mice was isolated and predicted loci with a CFD score□>□0.4 and an MIT score >□0.5 were PCR-amplified and Sanger sequenced. Animals without off-target mutations were used for generation of homozygous Plcγ2-P522R KI strain. Animals were raised and handled at the DZNE, Munich, and at the Laboratory Animal Center of the University of Eastern Finland, Kuopio. All animal experiments were carried out in accordance with the guidelines of the European Community Council Directives 86/609/EEC and approved by the Bavarian government and the Animal Experiment Board of Finland (ESAVI 21203-2019, EKS-004-2019).

### Mouse genotyping

Genomic DNA was purified from ear biopsies by isopropanol precipitation. The *Plc*γ*2* locus harboring the KI mutation was amplified by PCR using forward primer 5’-GCTGTCCTTCGGTGATGACA −3’ and reverse primer 5’-CAGACCGCCTGTTGGGAATA −3’. Resulting PCR products of 825 base pairs (bp) were further processed using the restriction enzyme StuI digesting the mutant KI allele into 412, 275, and 138bp fragments and WT to 412 and 413bp fragments. Fragments were subsequently detected from 1,5% agarose gel electrophoresis. The genotype of each mouse was verified by Sanger sequencing from the purified PCR product.

### Preparation of BMDM cultures

At the age of six months, KI male mice and their age- and gender-matched WT littermates were anesthetized with Ketamine-Xylazine mixture prior to trans-cardiac perfusion with ice cold saline (PBS). *Femur* and *tibia* bones were collected and processed for generation of BMDM cultures as described previously [10-12]. Briefly, bone marrow cells were flushed out using advanced RPMI 1640 medium (Life Technologies). Cells were differentiated for seven days in advanced RPMI 1640 supplemented with 2mM L-Glutamine, 10% (v/v)) heat-inactivated fetal calf serum (FCS), 100U/ml penicillin, 100μg/ml streptomycin and 50ng/ml macrophage colony stimulation factor 1 (m-CSF) (R&D System) in non-cell culture treated dishes. After seven days in culture, macrophages were carefully scraped, counted, and plated in optimal densities for subsequent analyses.

### Ip1 assay

Ip1 has been used as a standard readout when analyzing earlier identified *PLC*γ*2* variants in the previous studies [6, 13]. To measure Ip1 formation, 50000 KI (n=4) and WT (n=4) BMDMs per well were plated on a 96-well plate and allowed to attach overnight. IP-1 ELISA (Cisbio) assay was then used according to the manufacturer’s instructions. Some cells were treated with 25µM phospholipase C agonist, m-3M3FBS [7]. Ip1 concentration in each sample was calculated from the net optical density values using a standard curve. Values were normalized to the total protein concentration within each sample measured from the replicate wells. Data are normalized to the WT group and presented as mean ± SD.

### LDH cytotoxicity assay

For cytotoxicity assays, 10000 KI (n=4) and WT (n=3) BMDMs per 96-well plate well were seeded in RPMI 1640 (without phenol red) supplemented with 2mM L-Glutamine, 1% (v/v) heat-inactivated FCS, 100U/ml penicillin, and 100μg/ml streptomycin, and 50ng/ml m-CSF. After the cells were attached, the medium was replaced with m-CSF-depleted medium. Medium samples from triplicate wells per sample were collected 4h, 24h and 48h after m-CSF withdrawal. To determine maximum LDH release in each sample, replicate wells were treated with 2% Triton X-100 solution before medium collection. LDH cytotoxicity was measured according to the Cytotoxicity Detection Kit (LDH, Roche) protocol. LDH levels in each sample were normalized to the maximum LDH release in the corresponding sample. Data are normalized to the WT BMDMs and presented as mean ± SD.

### BMDM Phagocytosis assay

For phagocytosis assays, 10000-20000 KI (n=7) and WT (n=6) BMDMs were plated on a 96 well plate one day prior to the assay. The assay was repeated in two independent experiments. Phagozytosis was initiated by adding 5µg/well pHrodo-labeled bioparticles (pHrodo™ Red Zymosan BioParticles™ Conjugate for Phagocytosis, ThermoFisher Scientific, P35364) with or without 10 µM Cytochalasin D. Four fluorescent images per well with 300ms red channel acquisition time were taken every 15 minutes in total for three-hours using a 20x objective with the IncuCyte® S3 Live-Cell Analysis System (Sartorius). At the end of the assay, cells were incubated with 1µM Vybrant™ DyeCycle™ Green Stain (Invitrogen) for 0.5h and stained cell nuclei were imaged with 250ms green channel acquisition time for counting cells. Images were analyzed using the IncuCyte™ Analysis Software. pHrodo red fluorescence area averaged from four images taken per well were normalized to the corresponding cell count. Red fluorescent area was normalized to the WT group at the latest timepoint and shown as mean ± SEM.

### BV2 cell transfection and phagocytosis assay

pLenti-C-Myc-DDK (control) and human PLCγ2-myc-DDK (WT) in pLenti-C backbone vectors were obtained from OriGene (PS100064, RC200442L1). PLCγ2-myc-DDK was subjected to site-directed mutagenesis (QuikChange Lightning Multi Site-Directed Mutagenesis Kit, Agilent, 210515) to create PLCγ2-P522R-myc-DDK (P522R) constructs. Briefly, the pLenti-PLCγ2-myc-DDK plasmid was amplified using a single mutagenic primer (5’-AGTGCCCCAGGATATACGCCCTACAGAACTAC-3’). Subsequently, the original plasmid was digested with DpnI restriction enzyme and the remaining mutagenized plasmid was transformed into XL10-Gold® ultracompetent cells (Agilent). Integrity of the insert sequence and introduction of the point mutation were confirmed by Sanger sequencing of the isolated plasmid DNA.

For phagocytosis assays, BV2 microglia cells were transfected using Viromer Yellow kit (Lipocalyx) according to the manufacturer’s instructions with small adjustments. In brief, 50000 BV2 cells per well were seeded on a 12-well plate in RPMI 1640 (without phenol red) supplemented with 2mM L-Glutamine, 1% (v/v), fetal calf serum (FCS), 100U/ml penicillin, and 100μg/ml streptomycin one day prior to the transfection. One µg of WT, P522R, or control (CTRL) plasmids together with 0,32µl of Viromer Yellow in 100µl of Viromer Yellow buffer per well were used for transfections. Transfection medium was replaced four hours after transfections started. On the next day, cells were washed with PBS and incubated with pHrodo-labeled bioparticles (pHrodo™ Green E. coli BioParticles™ Conjugate for Phagocytosis, ThermoFisher Scientific) in OptiMem for three hours at +37°C. Cells were then washed with PBS, gently scraped in flow cytometry buffer (1% FCS, 2mM EDTA in PBS) and transferred to v-shaped 96-well plates. Non-viable cells were excluded by staining with 7-AAD Staining Solution (Abcam) for five minutes and pHrodo fluorescent signal was analyzed from 20000 live cells by flow cytometry. Fluorescent signals were normalized to unstained (no pHrodo-labeled bioparticles) control within each group. Phagocytic activity is shown as geometric median of the fluorescence intensity (MFI) and as % of phagocytic cells within the analyzed cell population as mean ± SD. MFI was normalized to the WT group.

Overexpression of the PLCγ2 WT and P522R plasmids was verified by Western blotting as described [37], using rabbit anti-PLCγ2 (1:1000, Cell Signaling Technologies) antibody, mouse anti-Myc (1:1000, Millipore) antibody, and mouse anti-β-actin (1:1,000, Abcam) antibody with respective secondary antibodies (1:5000, GE Healthcare).

### TNFα, IL-6, and IL-1β ELISA and NO assay

To address acute inflammatory responses, one million KI (n=4) and WT (n=3) BMDMs were plated on six-well plates and allowed to attach. Cells were then treated with 1µg/ml LPS and 20ng/ml IFN-γ in PBS or with vehicle (PBS) for three hours. Afterwards, media were collected and the levels of TNFα, IL-6, and IL-1β were determined using Mouse TNF alpha, IL-6, and IL-1 beta ELISA Ready-SET-Go!™ kits, respectively (Invitrogen™, eBioscience™). Cytokine levels were normalized to the total protein concentration in the lysate measured with BCA protein assay kit (Thermo Scientific). NO levels were analyzed using Griess Reagent Kit for Nitrite Determination (G-7921, Life Technologies) and normalized to the total protein concentration within the corresponding lysate. All kits were used as instructed by the manufacturers. Data are normalized to the WT BMDMs and presented as mean ± SD.

### NanoString gene expression analysis

Brains of the same six-month old KI and WT mice used for establishing the BMDM cultures were removed after transcardial perfusion and dissected into two hemispheres. Frozen right hemibrain was crunched in liquid nitrogen and 10-20mg of pulverized mouse brain was used for RNA extraction with RNeasy Plus Mini Kit (Qiagen). RNA concentration and quality were determined using Agilent RNA 6000 Nano Kit according to the manufacturer’s recommendations. Gene expression data of the KI (n=3) and WT (n=3) mice were generated with the mouse neuropathology gene expression panel of the *NanoString* Technologies using the nCounter system. Gene expression data were analyzed using nSolver Advanced analysis software (*NanoString* Technologies) with build-in quality control, normalization, and statistical analyses. Expression data of significantly altered genes for individual animals are shown as Log2-transformed fold changes. Themes annotated with individual genes are derived from the *NanoString* Neuropathology panel. Pathway analysis was done using pathway annotations provided by the panel.

### RT-qPCR analysis

For RT-qPCR analysis, RNA was reverse□transcribed into cDNA using the SuperScript III First□Strand Synthesis System (Thermo Fisher Scientific) according to the manufacturer’s protocol. Target specific PCR primers for mouse *Cst7* (5’-GTGAAGCCAGGATTCCCCAA-3’ and 5’-GCCTTTCACCACCTGTACCA-3), *Ccl3* (5’-CCAGCCAGGTGTCATTTTCC-3’ and 5’-AGTCCCTCGATGTGGCTACT-3’), and *Ctsd* (5’-AATCCCTCTGCGCAAGTTCA-3’ and 5’-CGCCATAGTACTGGGCATCC-3’) were obtained from TAG Copenhagen. FastStart Universal SYBR Green Master (Roche) was used for qPCR. The comparative ΔΔCt method was used to calculate *Gapdh* (5’-CAGGAGAGTGTTTCCTCGTCC-3 and 5’-TTCCCATTCTCGGCCTTGAC-3’) -normalized expression levels of the target mRNAs. Expression of *Tyrobp* (Mm.PT.58.6069426, IDT), *Clec7a* (Mm.PT.58.42049707, IDT), *and Plc*γ*2* (Mm01242530_m1, Thermo Fisher Scientific) were determined with mouse TaqMan assays and normalized to the expression of *Actb* (Mm.PT.39a.22214843.g, IDT) in the corresponding samples. Data are normalized to the expression levels in the WT group and shown as mean ± SD.

### Immunofluorescence analyses and confocal imaging

Following cardiac perfusion, left-brain hemispheres were immerse-fixed in 4% paraformaldehyde for 24h, followed by 30% sucrose for 24h. After freezing, 50μm microtome cut free floating sections were washed briefly and then blocked using 5% donkey serum for one hour at room temperature. Sections were then incubated with a Gfap antibody (1:500, Thermo Fisher Scientific) at 4°C overnight. Sections were washed and incubated in secondary antibody (donkey anti-rabbit 5551:1000, Thermo Fisher Scientific) for two hours at room temperature. Lastly, slides were washed and stained with 4′,6-Diamidin-2-phenylindol (DAPI, 5μg/ml) before mounting coverslips with ProlongTM Gold Antifade reagent (Thermo Fisher Scientific). Images were acquired using a LSM 710 confocal microscope (Zeiss) and the ZEN 2011 software package (black edition, Zeiss). Laser and detector settings were maintained constant for the acquisition. For analyses, at least three images were taken per slide using 10x (Plan-Apochromat 10x/0.45 M27) objective. For Gfap coverage analysis, confocal acquired images for cortex or hippocampus were imported to FIJI, and channels were separated by “Image/Color/Split Channels”. Following this, background noise was removed using Gaussian filtering and intensity distribution for each image was equalized using rolling ball algorithm. All layers from a single image stack were projected on a single slice by “Stack/Z projection”. Lastly, Gfap-positive staining was segmented using automatic thresholding method “Moments” in FIJI. Data are presented as mean ± SD.

### PET-imaging

For PET imaging, one-year-old KI (n=3) and WT (n=4) female mice were anesthetized with isoflurane (1.5% with N2/O2 70%/30% through nose cone). The mice were placed on a heated animal holder on the scanner bed in a prone position and secured with tape to prevent movement during scanning. The mice were imaged using a dedicated PET scanner (Inveon DPET, Siemens Healthcare) and immediately afterwards with CT (Flex SPECT/CT, Gamma Medica, Inc.) for anatomical reference images using the same animal holder. Dynamic imaging of 70 minutes was started at the time of the administration of the activity (18F-FEPPA: 15.8 ± 1.4 MBq and 18F-FDG: 13.9 ± 1.2 MBq) through the tail vein. Data were gathered in list-mode form, and corrected for dead-time, randoms, scatter and attenuation. Regions of interest (ROIs) were drawn for whole brain (excluding cerebellum and olfactory bulb), cerebellum, pons, hippocampus and frontal cortex using Carimas 2.10 software (Turku PET Centre, Finland). Also, for image derived input function (IDIF) a ROI was drawn for the *veca cava*. Logan analysis for 18F-FEPPA-data and Patlak analysis for 18F-FDG data was done using the IDIF to estimate the inflammation and cerebral glucose metabolism respectively for the brain ROIs.

### Statistical analyses

Statistical significance between groups was tested using independent samples *t-*test or Mann-Whitney U test depending whether the data fulfilled the assumptions for parametric tests or with two-way ANOVA (more than two groups) followed by LSD post-hoc test. Gene expression data were analyzed using nSolver Advanced analysis software (NanoString Technologies) with build-in quality control, normalization, and statistical analyses. Immunohistochemical data were checked for normality using the Shapiro-Wilk method, the D’Agostino and Pearson, as well as the Kolmogrov-Smirnov normality tests. Statistical significance was calculated using two-tailed unpaired *t*-test. All statistical analyses were performed using IBM SPSS Statistics 25 or GraphPad Prism software. A threshold for statistical significance was set at p<0.05.

## List of abbreviations

Plcγ2: phospholipase C gamma 2
Pip2: phosphatidylinositol 4,5-bisphosphate
Ip3: 1,4,5-trisphosphate
Dag: diacylglycerol
Trem2: Triggering receptor expressed on myeloid cells 2
Nfκb: nuclear factor kappa-light-chain-enhancer of activated B cells
Mapk/Erk: mitogen-activated protein kinase
Akt: protein kinase B
KI: knock-in
WT: wild type
BMDM: bone marrow-derived macrophage
Ip1: inositol monophosphate
LPS: lipopolysaccharide
IFNγ: interferon gamma
TNFα: tumor necrosis factor alpha
IL-6: interleukin-6
IL-1β: interleukin 1 beta
NO: nitric oxide
DAM: disease associated microglia

## Declarations

### Ethics approval and consent to participate

Animals were raised and handled at the DZNE, Munich, and at the Laboratory Animal Center of the University of Eastern Finland, Kuopio. All animal experiments were carried out in accordance with the guidelines of the European Community Council Directives 86/609/EEC and approved by the Bavarian government and the Animal Experiment Board of Finland (ESAVI 21203-2019, EKS-004-2019).

### Consent for publication

Not applicable

### Availability of data and materials

All data generated or analysed during this study are included in this published article and its supplementary information files.

### Authors’ contributions

MT, MH and CH designed the study, interpreted the data and wrote the paper. MT also designed experiments, collected and processed cell and brain samples, generated functional and RNA data, performed related analyses, and generated the figures. RW generated PLCγ2 variant plasmids and RW and HM performed phagocytosis assays of the BMDM cells. BW designed and generated the knock-in mouse line. SP performed immunohistochemical staining and related analyses. TK assisted in the processing of the NanoString data. KJ performed a FEPPA and FDG-PET imaging and related data analysis for one-year-old mice. PP produced FEPPA and FDG ligands for PET imaging. XX helped with processing the RNA samples. PM assisted with the sample collection and ELISA assays. MM and AH assisted with the data interpretation and revised the manuscript. All authors read and approved the final manuscript.

## Acknowledgements

We thank Mrs. Maarit Pulkkinen for her expert technical assistance with PET imaging.

