## Supplemental figures for "The Alzheimer’s disease-associated protective *Plcγ2*-P522R variant promotes beneficial microglial functions"

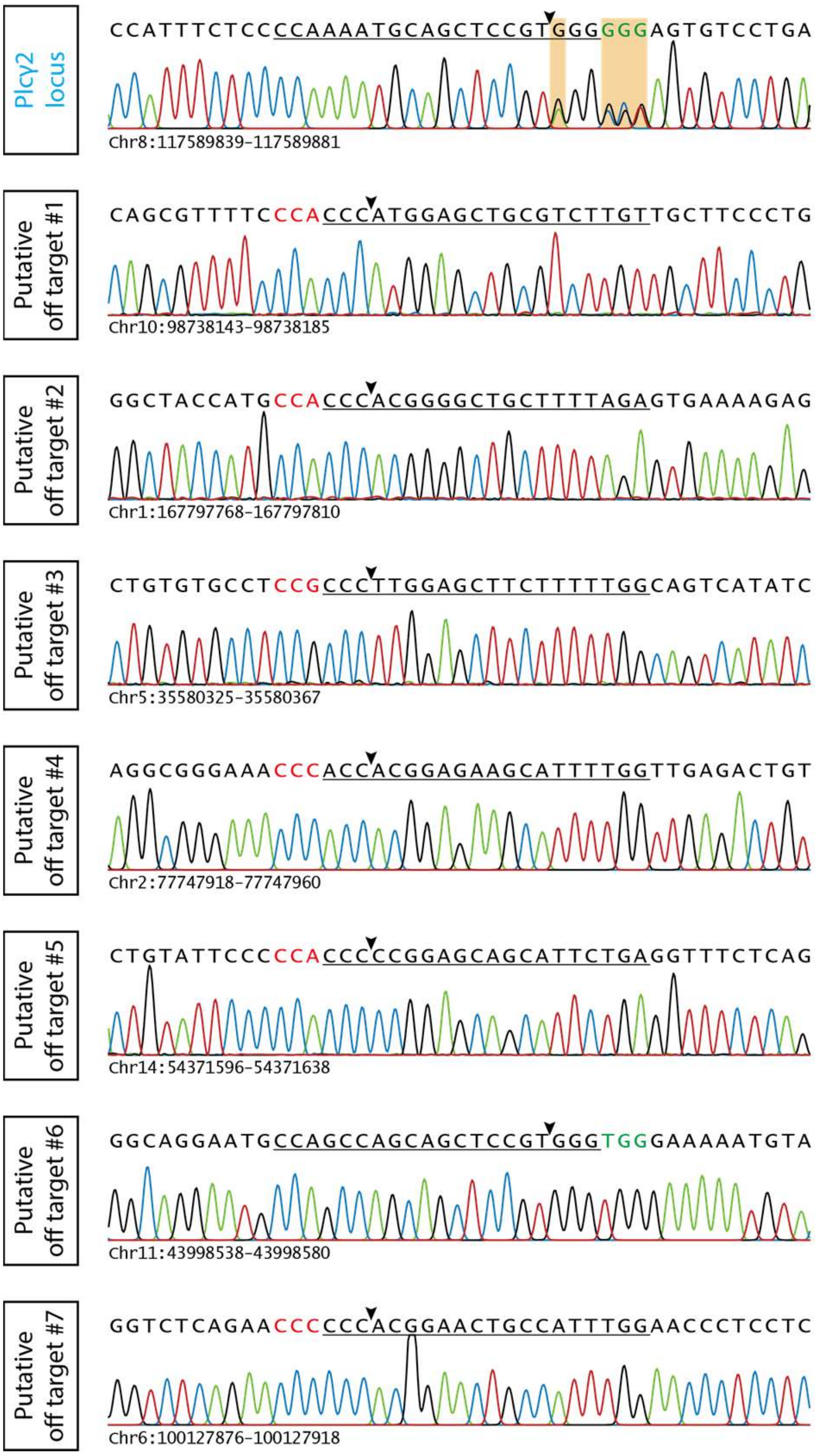


**Supplementary figure 1:** Representative Sanger-sequencing chromatograms of the Plcγ2 on-target site and the seven putative off target sites of a heterozygous F1 animal. Mixed peaks in the Plcγ2 locus show the correct P522R substitution (CCC>AGG, on complementary strand) and a silent mutation for genotyping purposes. Underlined: Protospacer; arrow head: putative cut site; green letters: PAM site on shown strand; red letters: PAM site on complementary strand.

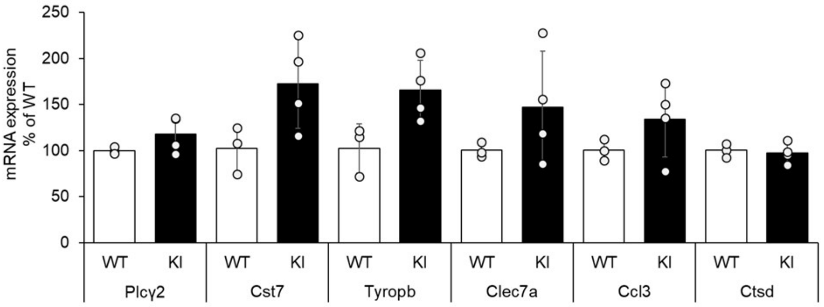


**Supplementary figure 2:** RNA expression levels of Plcγ2 and microglia specific disease-associated microglia signature genes, Cst7, Tyropb, Clec7a, Ccl3, and Ctsd in the brain of six-month-old Plcγ2-P522R homozygous knock-in (KI) and wild type (WT) mice. Normalized to the WT gourp, mean ± SD, n=3-4 per genotype.
